# Computer simulation of knotted proteins unfold and translocation through nano-pores

**DOI:** 10.1101/378901

**Authors:** M. A. Shahzad

## Abstract

We study the unfold and translocation of knotted protein, YibK and YbeA, through α-hemolysin nano-pore via a coarse grained computational model. We observe that knot of protein unfold in advance before the translocation take place. We also characterized the translocation mechanism by studying the thermodynamical and kinetic properties of the process. In particular, we study the average of translocation time, and the translocation probability as a function of pulling force *F* acting in the channel. In limit of low pulling inward constant force acting along the axis of the pore, the YibK knotted protein takes longer average translocation time as compare to YbeA knotted protein.

## I. INTRODUCTION

The passage of a molecule through a biological pore, the so-called translocation, is a fundamental process occurring in a variety of biological processes [1], such as virus infection [2], and gene expression in eukaryotic cells [3]. The translocation process involves the exchange of proteins, ions, energy sources, genetic information and any particle or aggregate that plays a role in cell functioning and living. For instance, translocation of biopolymer across membranes channel is universal in biology and includes the passage of RNA strands inside nuclear pores [4], absorption of oligonucleotides on suitable sites [5], transport of protein back and forth from the cells [6], mitochondrial import, protein degradation by Adenosine triphosphate (ATP)-dependent proteases and protein synthesis [7, 8].

The biological cell is covered by all types of pores and channels that control exchange of ions and molecules between sub-cellular compartment. These pathways are of vital importance to cellular function. Examples are: ion channels at the cell surface that control the flow of ions; pores inserted into cell membranes upon viral infection that serve as conduits for genome transfer; the nuclear pore complex (NPC) that regulates the transport of mRNA and proteins across the nuclear envelope of eukaryotic cells; and pore across the used for protein secretion into cell organelles.

Proteins play an important role in transport process across the cell membranes. The transport can be either passive or active. The passive transport involves the movement of biochemicals from areas of high concentration to areas of low concentration and requires no energy for transportation. The passive transport can take place by diffusion or by interposition of carriers to which the transported substances are bounded. On the other hand, active transport are biological processes that move oxygen, water and nutrients into cells and remove waste products. Active transport requires chemical energy because it is the movement of biochemicals from areas of lower concentration to areas of higher concentration. In active transport, trans-membrane proteins use the Adenosine-Tri-Phosphate (ATP) chemical energy to drive a molecule against its concentration gradient. A mechanism of selection, based on the interaction between signal recognition particles and relative receptors, allows the trans-membrane proteins to detect the system to be transported.

Proteins are not only the building blocks of transmembrane channels, but are also commonly transported between organelles and cells, in folded and unfolded form. Very often membrane apertures are not large enough to enter the proteins in their native (folded) conformations. For instance, the narrowest constriction in the proteasome organelle is only 13 Å in diameter. It is thus necessary for proteins to unfold in advance or during their passage across the pore [10, 11].

Here we used a coarse-grained computational model to simulate numerically the unfold and translocation process for the knotted protein through a *α*-hemolysin channel. We show that the knot in knotted protein unfold during the translocation process.

## II. COMPUTER MODELING OF STRUCTURED POLYPEPTIDE CHAIN AND NANOPORE

We used a coarse-grained computational model to simulate numerically the translocation process for the protein through a channel. We consider the Go-like model as a natural approach to access the impact of the molecules structural properties on translocation. Below we show in detail the importing mechanism, pore and particle interaction potential, numerical integration scheme and equation of motion for the material points constituting the system.

### A. Protein numerical model: The Gō-like approach

Protein folding is a complex process involving large number of degree of freedom. A linear polypeptide protein chain is organized into a space-filling, compact, and well-defined 3-dimensional structure. The monomeric unit in a polypeptide chain is the peptide group. The C,O,N,and H atoms in the amino acid lie in the same plane, and successive planes defined angels *ϕ* and *ψ*. The conformation of a chain of *n* amino acids can be defined by 2*n* parameters. Therefore, performing traditional molecular dynamics simulation with atomistic detail poses a challenge if one wished to exhaustively explore the folding landscape of a moderately sized protein with finite computation resources. For this reason coarse-grained representation of the polypeptide chain have been employed in folding simulation. The phenomenological off-grid model proposed by Nobuhiro-Go and Harold A. Scheraga [12, 14] is a coarse-grained model that identifies all the amino acids with their C*α* carbon atoms. Rather than representing each atom in the protein explicitly groups of atoms can be treated as a single coarse-grained site. This way, each residue corresponds to a single coarse-grained site placed at the C*α* carbon atom. Through this way, protein are reduced to a chain of material points coinciding spatially with the C*α* carbon atoms of the protein backbone and no side chain characteristic are retained. The atomic mass of the material points are the same and equal to 1 in the so-called code measure units or internal units. Different refinements of this model are available in literature [13, 15–19], and are generally referred as Go-Like models. The approach presented in our numerical work is adopted from reference [13, 20, 22, 23].

The native structure of a protein is taken from the protein data bank (PDB) file and the interaction between the amino acids are grouped into native and non-native state. The protein data bank (PDB) format file contains information about atomic coordinates for various types of proteins, small molecules, ions and water. It provides a standard representation for macromolecular structure data which is derived from X-ray diffraction and nuclear magnetic resonance (NMR) studies.

In Gō-like model, energy minima are assigned to a particular reference configuration, usually coincident with the native structure of the considered protein. This is why Go-models, where only native contacts contribute to the free energy, can be properly applied to small globular protein whose crystalline structure is already know and the folding dynamics are mainly determined by the topology of the natured conformation. The Gō-like model generates almost ideal folding pathway. Also, the native conformation so obtained are stable and closed to the reference one, which can be extracted from PDB. The Gō-like model is ideal for two-state folding dynamics simulation, but it has been used also to predict intermediate configuration [13].

The main limitation of the Go-model is that its topological nature does not account for chemical factors, determining a considerable disagreement between real and simulated time scale. However, these discrepancies do not alter the folding thermodynamics and the correct sequence of events. In most cases the model allows the correct assessment of the free energy and of folding pathways.

Using the Gō-like approach for the numerical simulation, we used the following potentials acting on the material point of the protein: (1) Peptide potential (or bond potential) *V_p_*, (2) Bending angle potential *V_θ_*, (3) Twisted angle potential *V_ϕ_*, and (4) Non-bonded interactions (Lennard-Jones potential or barrier) *V_nb_*. These interactions are the simplest potential energy function that can reproduce the basic feature of protein energy landscapes at a mesoscopic level of detail, and it has proved to give insight into a remarkably broad range of properties. The combinations of a potential energy function and all the parameters that go into it constitutes a so-called force field.

Let ***r***_*i*_(*i* = 1,…,*m*) be the position vector of the m residues identified by their C*α* Carbon atom in the reference (native) configuration, and 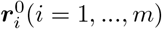 be the position vector of m residue in the current configuration. The peptide potential, responsible for the covalent bonds between the beads of the polymer chain, has the following expression:

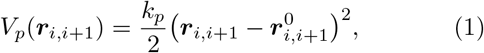

where ***r***_*i,i*+1_ = ***r***_*i*_ – ***r***_*i*+1_, 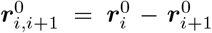 are the position vector between the bonded residues *i* and *i* + 1 in the instantaneous and native configuration respectively. The norm of position vector is the bond length. The empirical constant 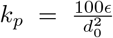 is in term of the equilibrium length parameter *d*_0_, and is in the unit of energy parameter *ϵ*. In our simulation, *d*_0_ = 3.8 Å is the average distance between two adjacent amino acids and *ϵ* sets the energy scale of the model. The bond potential is nothing more than a simple harmonic potential with spring constant *k_p_*.

The angular potential *V_θ_* is used to recover the secondary structure of protein in reference native conformation. Mathematically, it is equivalent to peptide potential *V_p_* by replacing the relative displacement by angular difference, that is

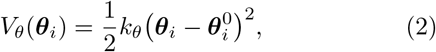

where *k_θ_* = 20*ϵ* rad^−2^ is the elastic constant expressed in term of the energy computational unit *ϵ*, and *θ_i_*, 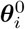 are bond angles formed by three adjacent beads in the simulated (time-dependent) and native conformation, respectively.

The dihedral potential (torsion) is 1-4 interaction, and are expressed as a function of the twisted angles *ϕ_i_* and 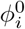, again refereed respectively to the actual and crystal configuration. The dihedral potential is important for the recovery of the correct protein secondary structure. The twisted angle is the angle formed between the two planes determined by four consecutive amino acids along the chain. The definition of twisted angle potential *V_ϕ_* is

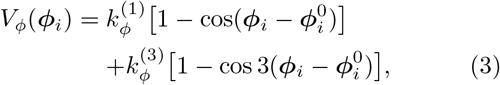

where 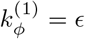 and 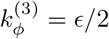 are dihedral constants expressed in term of energy unit *ϵ*.

Non-bonded (nb) interactions between nonconsecutive amino acids are modeled with Lennard-John 12-10 potential. In Gō-like model a distinction is made among the pairs of residues that interact following a potential that has also an attractive part, in addition to a repulsive part. The criteria for this distinction is made on the basis of the native distance with respect to a parameter of the model, the so-called cut-off radius, *R_c_*. Native contact [24] are identified by the cut-off distance *R_c_* to be chosen such that two residues *i* and *j* form a native interaction if their distance *r_ij_* in the native state is less than cut-off distance *R_c_*. This criterion selects a certain number of native contacts on the Proteins crystallographic structure. On the other hand, when two residues are not in native contact *r_ij_* > *R_c_*, they interact only through the Lennard-Jones repulsive tail (*σ/r_ij_*)^12^, where *σ* = 4.5 Å is a free length parameter correlated with the extension of the excluded volume (self-avoiding polymer chain). In other words, such residues in the protein chain will interact attractively if brought to a distance greater than the native one and repulsive otherwise. The expression for Lennard-Jones potential is:

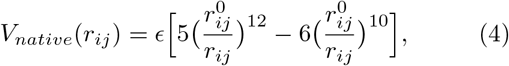

where all the symbols have been already defined. When 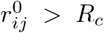, the purely repulsive contribution *V_nonnative_* is assigned to the pair of amino acids considered. This term enhances the cooperativity in the folding process and takes the form of a Lennard-Jones barrier

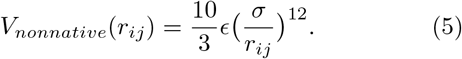

The non-bonded potential *V_nb_* summarized the possible long range interaction just described above and reads as

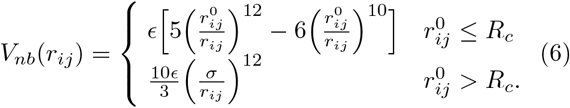

The total potential acting on all the residues of the proteins is then:

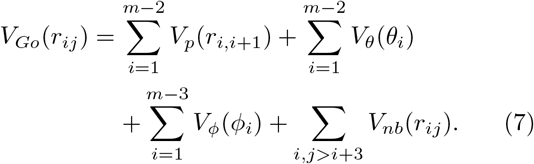

Figure (1, left upper Panel) illustrate the basic atomic structure of YibK knotted protein. The secondary structure with beta-sheets and *α*-helices is shown in fig (1, right upper Panel). The coarse-grained representation of knotted protein is described in fig (1, left lower Panel) with color that vary according to its residues type. Figure (1, right lower Panel) illustrate the Yibk knotted protein with trefoil knot region marked my rectangular. The detail structure of YbeA knotted protein are shown in Figure (2 (Figures produced by VMD-software).

**FIG. 1.**
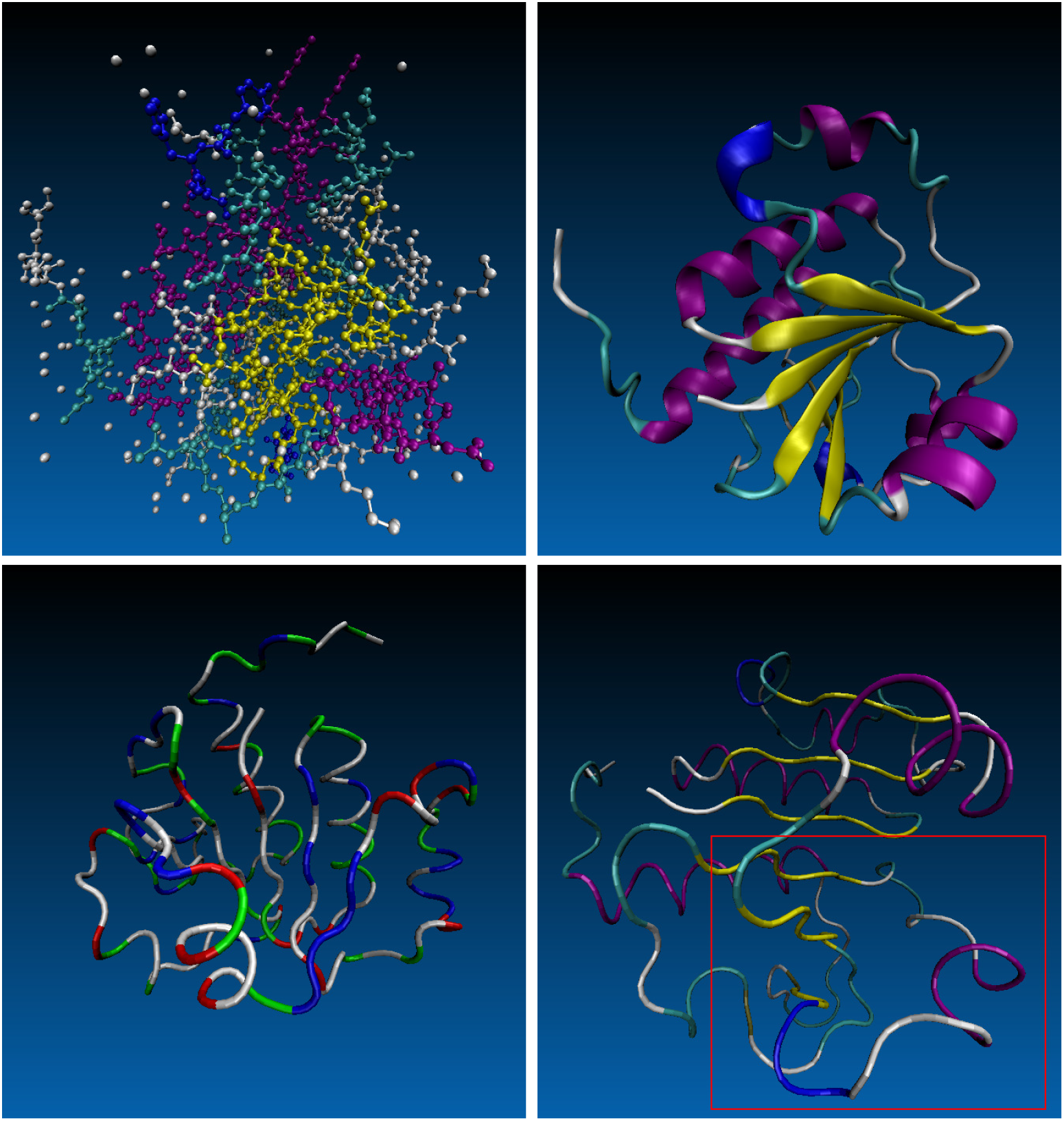
Left upper panel: Full atomic representation of YibK knotted protein. Right upper panel: Secondary structure of YibK revealed from its protein data band file, PDB-ID: IJ85. Lower left panel: YibK protein coarse grained description. In Go model only the *C_α_* carbon atoms along the main backbone chain of the protein are depicted. Lower right panel: The rectangle represent the knotted region.

**FIG. 2.**
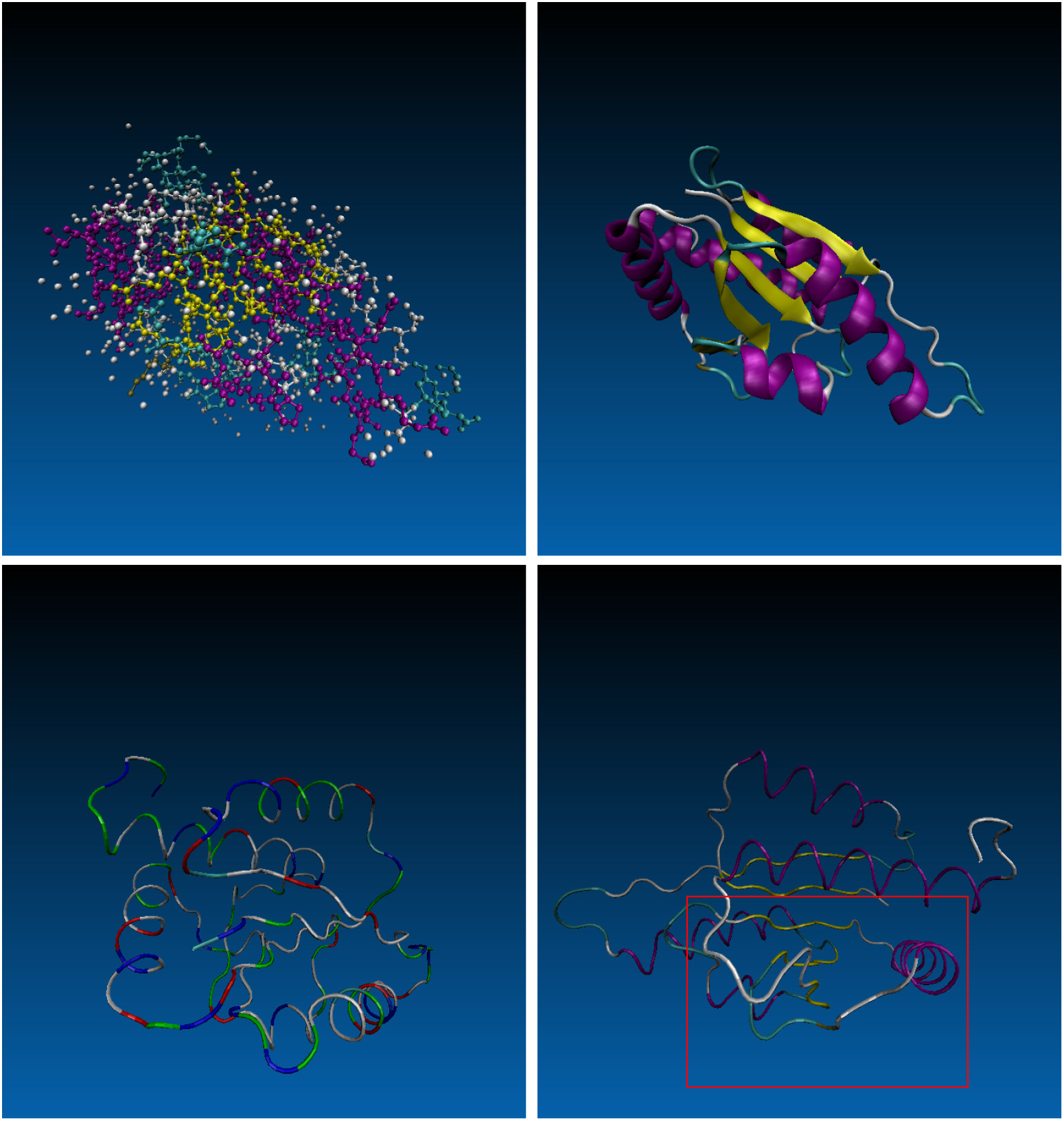
Structure of YbeA from Escherichia. Left upper panel: Full atomic representation of YbeA knotted protein. Right upper panel: Secondary structure of YbeA revealed from its protein data band file, PDB-ID: 1NS5. Left lower panel: YbeA protein coarse grained description. In Gō model only the *C_α_* carbon atoms along the main backbone chain of the protein are depicted. Right lower panel: The rectangle represent the knotted region.

### B. Pore Model

The confinement effect on protein dynamics can be represented by a step-like soft-core repulsive cylindrical potential. The cylinder axis of symmetry is set adjacent with the *x*-axis of the frame of reference used for protein translocation simulation. The same *x*-axis direction is used to develop the mechanical pulling of the protein by using a constant force *F_x_* applied to the foremost beads inside the confinement. The constant force is used in analogy with the electrical potential in voltage driven translocation experiment. The schematic representation of the system, the potential and the pulling mechanism are shown in Fig. (3).

The expression of the pore potential is given by:

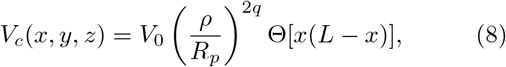

where *V*_0_ = 2*ϵ* and Θ(*s*) = [1 + tanh(*αs*)]/2 is a smooth step-like function limiting the action of the pore potential in the effective region [0, *L*]. *L* and *R_p_* are pore length and radius respectively. Also, 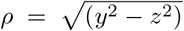 is the radial coordinate. The parameter *q* tunes the potential (soft-wall) stiffness, and *α* modulates the soft step-like profile in the *x*-direction; the larger the *α*, the steeper the step. In our simulation, we consider *q* = 1 and *α* = 2 Å^2^. The driving force *F_x_* acts only in the region in front of the pore mouth *x* ∈ [−2, 0], and inside the channel [0, *L*]. Pore length *L* = 100 Å and radius *R_p_* = 10 Å are taken from *α*HL structure data [25].

**FIG. 3.**
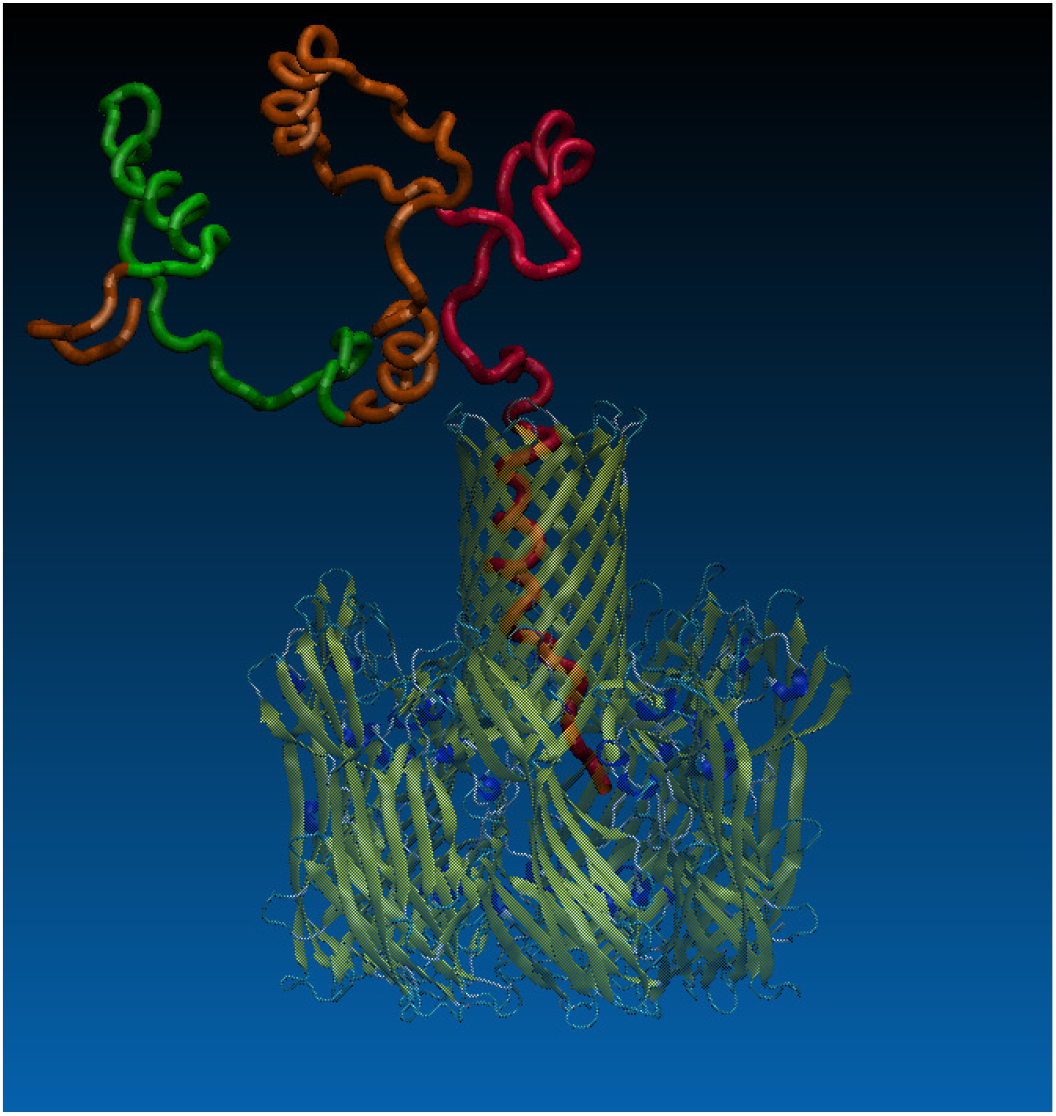
Schematic view of knotted protein pulled across a *α*-hemolysin nano-pore from CIS to TRANS-side. The confinement effect inside the pore is model by a step-like [0, *L*] cylindrical soft-core repulsive potential. A constant force *F_x_* in the direction of *x*-axis is used to drive the protein inside the channel. The force is always applied to the foremost residue inside the pore.

### C. Equation of Motion: Langevin dynamics

The equation of motion which governs the motion of material point is the well-know Langevin equation which is usually employed in coarse-grained molecular dynamics simulation. Consequently, numerical investigations are performed at constant temperature. The over-damped limit is actually implemented 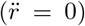 and a standard Verlet algorithm is used as numerical scheme for time integration [26]. The Langevin equation is given by

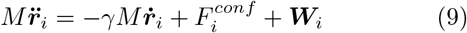

where 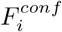 is the sum of all the internal and external forces acting on residue *i*. Here *γ* is the friction coefficient used to keep the temperature constant (also referred as Langevin thermostat). The random force ***W***_*i*_ accounts for thermal fluctuation, being a delta-correlated stationary and standard Gaussian process (white noise) with variance 〈*W_α_*(*t*)*W_α_*(*t*′)) = 2*γMRTδ_α,α′_δ*(*t* – *t*′). The random force satisfies the fluctuation-dissipation theory; the mean-square of *W* is proportional to the corresponding friction coefficient *γ*.

## III. RESULTS AND DISCUSSION

We used Langevin molecular dynamics simulation (Verlet integration scheme) to transport knotted protein YbeA (PDB ID 1NS5) and YibK (PDB ID IJ85) across a static *α*-hemolysin pore. The folding temperature occurred at *T*\* = 0.77 in reduced temperature units *R/ϵ*, corresponding to the experimental denaturation temperature *T* = 338 K. This defined the energy scale to the value *ϵ* ≃ 0.88 kcal mol^−1^. Using the time scale *t_u_* = *σ*(*M*/120*ϵ*)^1/2^ we can obtained the physical time unit from the simulation time. With *ϵ* ≃ 0.88 kcal mol^−1^, *σ* = 4.5 Å, and assuming the average mass of amino acid is *M* ~ 136 Da, we get *t_u_* ~ 0.25*ps*. In computer simulation the time step and friction coefficient used in the equation of motion (Langevin dynamics) are *h* = 0.001*t_u_* and 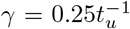, respectively. Further, the unit of force is defined as *f_u_* = *ϵ* Å^−1^ ~ 6 pN.

### A. Knotted protein Unfold

The protein in its native conformation is placed initially at the CIS-side of the pore. As shown in fig (4, upper left and right panel) the knotted region of Yibk knotted protein are marked by rectangle with residues color follow by red, blue, green, yellow, black and white. During the first simulation run time the knot unfold as describe in figure () with the same residue colors. In contrast to Piotz Szymczak [27] finding where an on/off pulling force untie the knot of protein, we observed that a simple constant pulling force can able to unfold the knot of knotted protein. Figure (4, lower left and right panel) show the initial fold conformation of YbeA knotted protein at the begiing of simulation. The lower right panel of figure () decribe its unfold conformation. The mechanism of unfold of the trefoil knot in our simulation is similar to fold mechanism as observed by Stefan Wallin *et al*. [28].

**FIG. 4.**
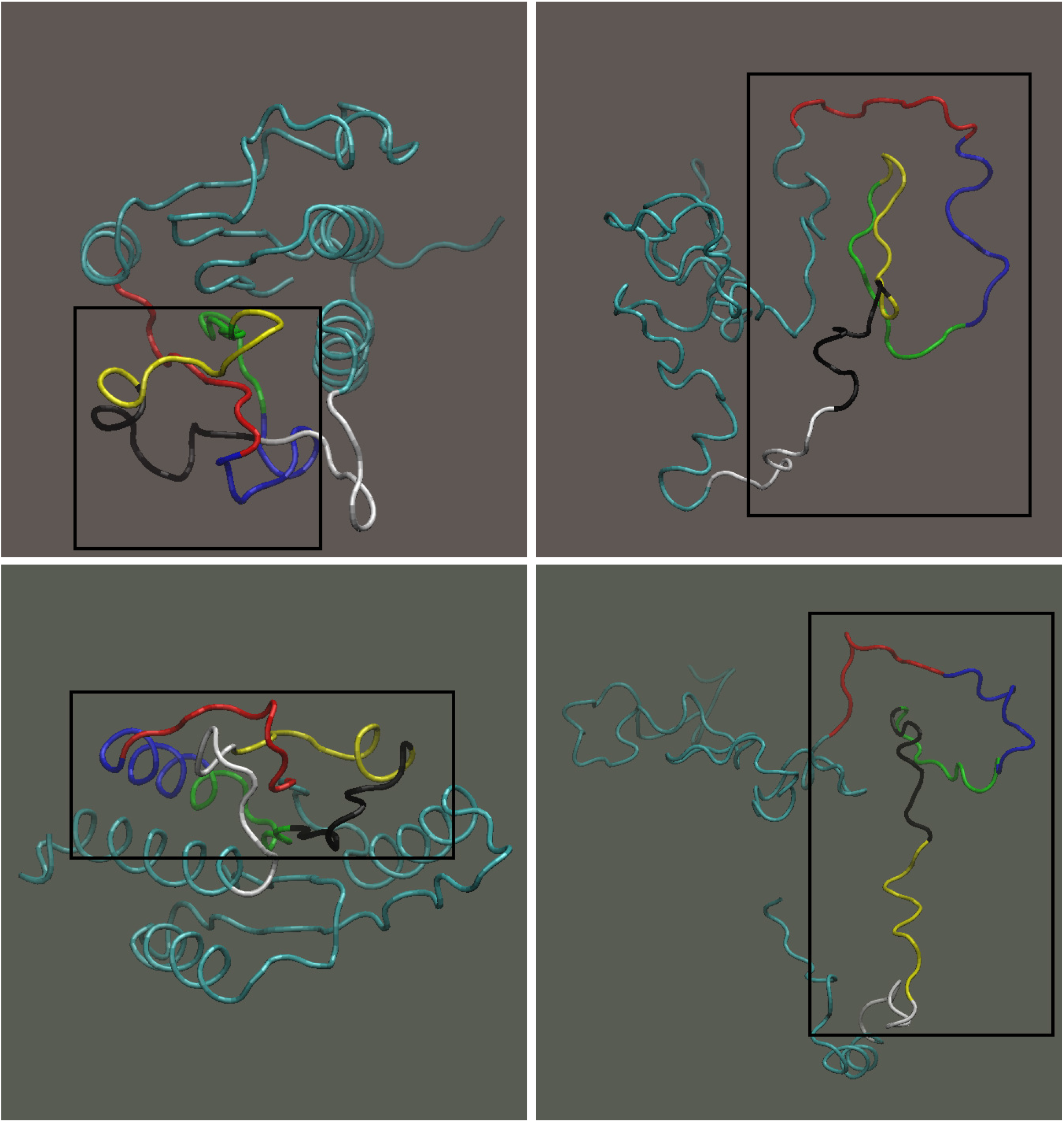
Snapshots of unfold mechanism. Upper left Panel: YibK protein with knot region marked by rectangle. Upper Right Panel: Unfold of the knot during the first simulation time. Lower left Panel: YbeA protein with knot describe by rectangle. Lower Right Panel: Unfold of the knot obtained after the first simulation run time.

### B. Knotted protein translocation

The simulation was run until the protein was fully expelled out from the cis to tans-side, and almost complete refolding occurred. However, the translocation may fail at low external force *F* within a large assigned waiting time *t_w_*; in this case, the run stops, discards its statistics, and restarts a new trajectory. Figure (5) illustrates the plot of translocation probability *P_Tr_* as a function of force. The probability of translocation *P_Tr_* can be defined as the number of translocation successes over the number of total runs, within the time *t_w_* ~ 15^3^*t_u_*. The translocation becomes (*P_Tr_* > 1/2) when the force exceeds a critical value *F_c_* ≃ 0.75*f_u_* for YbeA and *F_c_* ≃ 0.5*f_u_* for YibK knotted protein, while below these thresholds the probability goes rapidly to 0. The red square data represent the translocation probability for YibK protein while the filled blue circle describe the YbeA knotted protein translocation probability data.

Defining the collective variable coordinate *Q*,

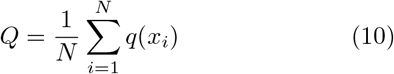

where *x_i_* is the *x*-coordinate of the *i*th bead and *q*(*x*) is the piecewise function defined as

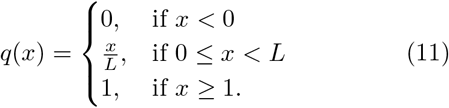

**FIG. 5.**
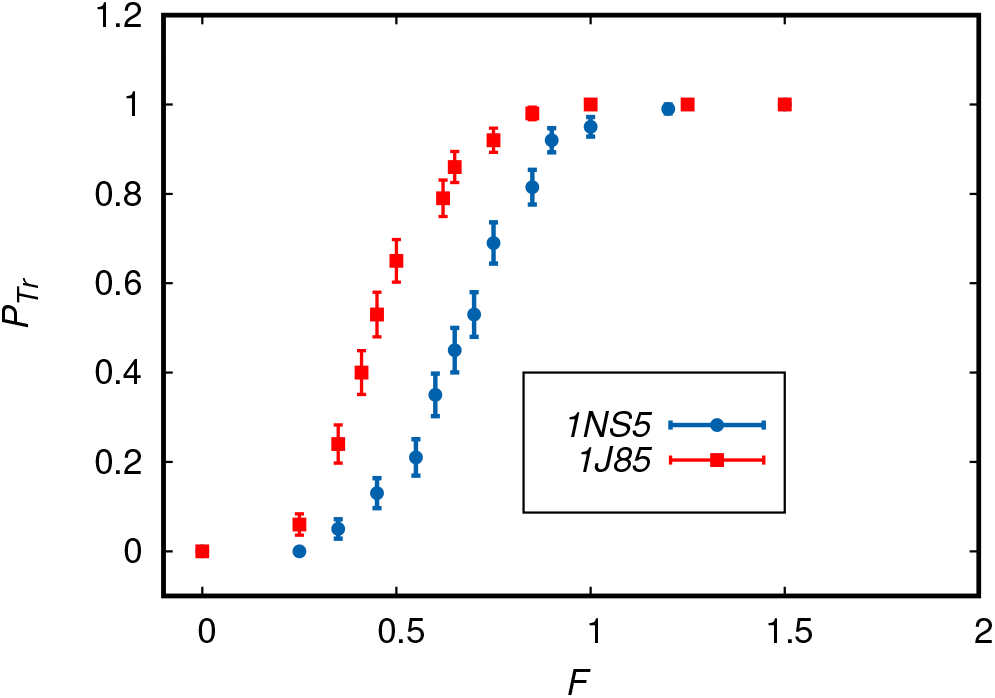
Ubiquitin translocation probability *P_Tr_* as a function of force. The error-bar are calculated using the statistical error formula.

The value of collective variable coordinate *Q* = 0 corresponds to the case when the folded protein is completely on the Cis-side of the nano-pore, and *Q* = 1 when the protein is completely translocate on Trans-side. The value 0 < *Q* < 1 represent the coordinate when the beads is inside the nano-pore. Figure (6) and (6) illustrate the few trajectory of knotted protein in collective variable coordinate *Q* as a function of time during the unfolding and translocation through a *α*-hemolysin pore.

**FIG. 6.**
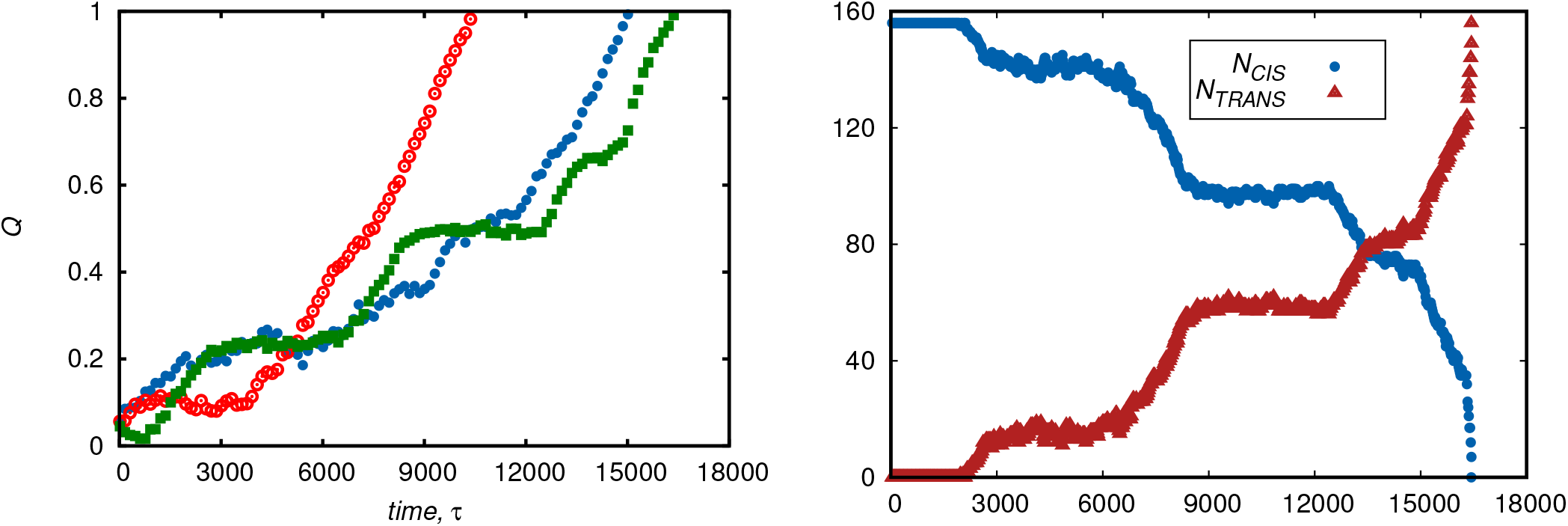
Trajectory of YibK knotted protein in collective coordinate *Q* as a function of time using a constant external pulling force of *F* = 0.75 at the C-terminus. YibK: A trace of the number of peptide count at the CIS-side (filled blue circle), and TRANS end (filled brown triangle).

**FIG. 7.**
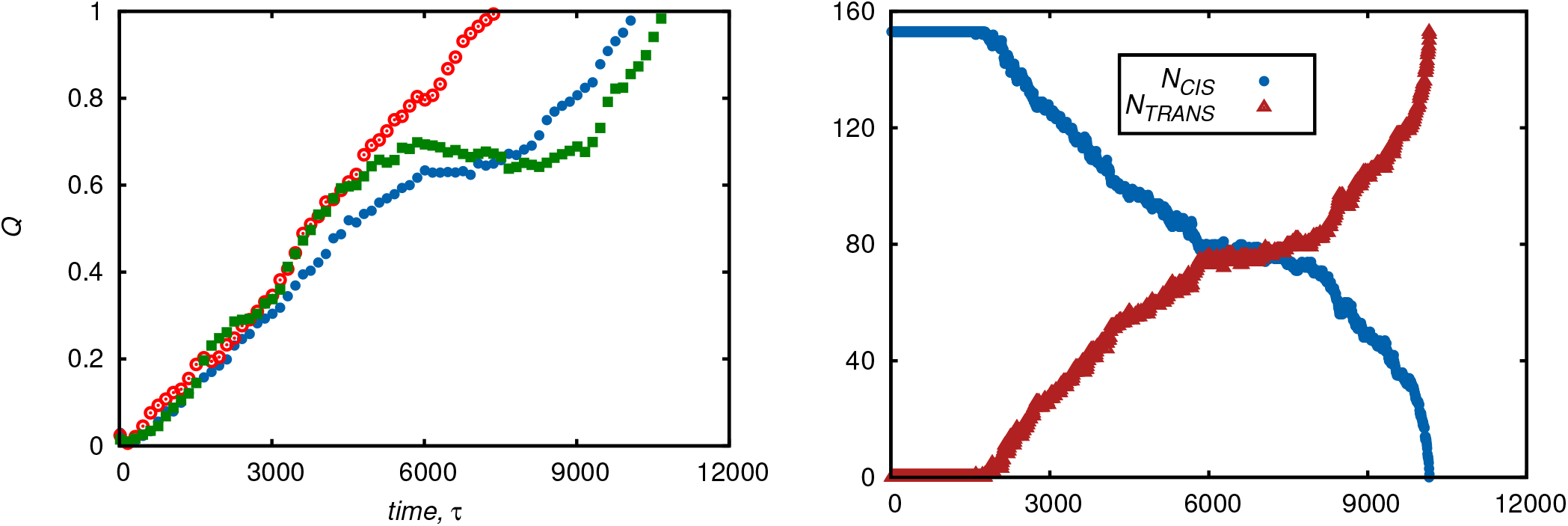
Trajectory of YbeA knotted protein in collective coordinate *Q* as a function of time using a constant external pulling force of *F* = 0.75 at the C-terminus. YbeA: A trace of the number of peptide count at the CIS-side (filled blue circle), and TRANS end (filled brown triangle).

**FIG. 8.**
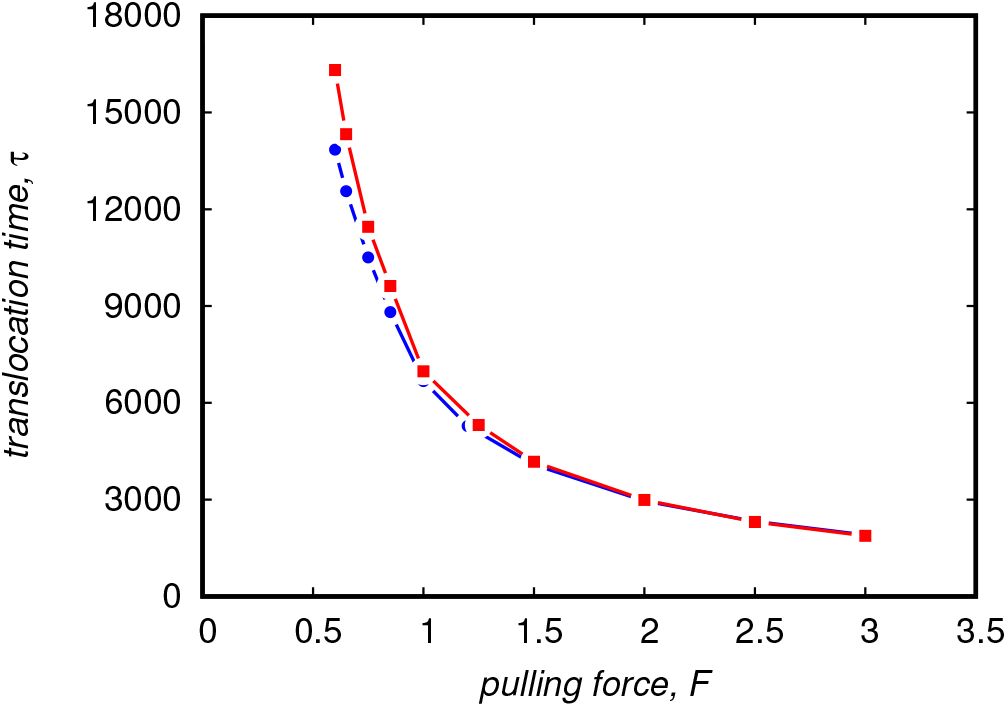
Average translocation time *τ* of ubiquitin as a function of the force *F*. The red square circle represent YibK average translocation time while the blue circle show the YbeA translocation time data.

The translocation process can be further characterized by studying the statistics of translocation times. The time statistics of translocation events is accessible to experiment [11] in which current loop drops signal the occupation of the channel by the passing molecule; for this reason, these times are also called the blockage times. In the simulation, the translocation time can be measured as first arrival time *t* at the channel end *x* = *L* of the protein center of mass. The average translocation time is shown in Fig. (8) as a function of force *F*.

## IV. CONCLUSION

We have investigated the mechanical unfold and translocation of knotted protein through a biological *α*-hemolysin nano-pore using coarse-grained computational model. In this simplified model translocation process we import a knotted protein via a finite size channel by a uniform pulling force acting inside of the channel only. In particular, we present in detail the mechanical unfold and translocation of trefoil knotted protein Yibk and YieA. Our finding shows that the knot of the protein unfold in advance before or during the translocation process. We examine the kinetic and thermodynamical characteristic of the process by studying the statistics of blockage times and the translocation probability as a function of the pulling force *F* acting in the pore. As shown in the Fig. (5) of probability of the translocation as a function of force, the translocation dynamics occurs when the importing force exceeds a threshold value *F_c_* depending on the free-energy barrier that protein has to overcome in order to slide along the channel. Such a free-energy barrier results from competition of the unfolding energy and the entropy associated with the confinement effects of the pore. Mathematically, the molecular dynamics simulations at the coarse-grained level of protein translocation can be described by a simple drift-diffusion Smoluchowski equation exploiting the first passage theory and radiation boundary condition.

## Supporting Information

**Movie S1** This movie file (produced by VMD software) represents the unfold and translocation of **YbiaK** through *α*HL nanopore having length *L* = 100 Å and radius *R_p_* = 10 Å.

**Movie S2** This movie file (produced by VMD software) describes the unfold and translocation of **YbeA** through *α*HL nanopore having length *L* = 100 Å and radius *R_p_* = 10 Å.

